# A hybrid method for discovering interferon-gamma inducing peptides in human and mouse

**DOI:** 10.1101/2023.02.02.526919

**Authors:** Anjali Dhall, Sumeet Patiyal, Gajendra P. S. Raghava

## Abstract

A host-specific technique has been developed for annotating interferon-gamma (IFN-γ) inducing peptides, it is an updated version of IFNepitope. In this study, dataset used for developing prediction method contain experimentally validated 25492 and 7983 IFN-γ inducing peptides in human and mouse host, respectively. In initial phase, machine learning techniques have been exploited to develop classification model using wide range of peptide features. In most of the case, models based on extra tree perform better than other machine learning techniques. In case of peptide features, compositional feature particularly dipeptide composition performs better than one-hot encoding or binary profile. Our best machine learning based models achieved AUROC 0.89 and 0.83 for human and mouse host, respectively. In order to improve machine learning based models or alignment free models, we explore potential of similarity-based technique BLAST. Finally, a hybrid model has been developed that combine best machine learning based model with BLAST and achieved AUROC 0.90 and 0.85 for human and mouse host, respectively. All models have been evaluated on an independent/validation dataset not used for training or testing these models. Newly developed method performs better than existing method on independent dataset. The major objective of this study is to predict, design and scan IFN-γ inducing peptides, thus server/software have been developed (https://webs.iiitd.edu.in/raghava/ifnepitope2/).

**Highlights:**

- An updated method for predicting interferon-gamma (IFN-γ) inducing peptides.
- A wide range of features have been generated using Pfeature tool.
- Models were trained and tested on experimentally validated datasets.
- Hybrid models developed by combining machine-learning and BLAST.
- IFNepitope2 server is available to design subunit or peptide-based vaccines.

## Introduction

Type II interferons or IFN-γ is a multifunctional pleiotropic cytokine play an essential role in innate and acquired immune responses. IFN-γ is majorly secreted by helper T cells, cytotoxic T cells, natural killer cells, B cells, macrophages [1–5]. Studies report that, IFN-γ possess both pro-inflammatory and anti-inflammatory properties for example it upregulates MHC-I/II protein expression, modulates the production of T helper cells (Th1), activates tumoricidal activity and inhibit the expression of immunosuppressive cytokine (IL-17) [6–8]. Moreover, it is involved in intracellular communication, tumour cell identification and eradication [1, 9]. Recent studies report that, interferons can act as the potential therapeutic candidates in various infectious diseases and cancer [10, 11]. For example, Yan et al., shows that IFN-γ acts as a prognostic marker for PD-1/PD-L1 immune checkpoint inhibitor cancer therapy and associated with the better response to immune checkpoint blockade (ICB) therapy in cancer patients [12, 13].

Peptide-based cancer immunotherapies have great potential to treat a variety of malignant tumours [14]. Peptides acts as versatile immune vaccine candidates that stimulate both innate and adaptive immune system by interacting with various immune cells such as dendritic cells, NK cells, T cells and B cells [15–18]. In order to generate anti-tumor or anti-viral immune response, the potential peptide-based vaccine candidate should activate the production of Th1 associated cytokines such as IFN-γ, IL-2, IL-12, and TNF-α, and inhibit the production of Th2 cytokines including IL-4, IL-6 and IL-13 [19]. Studies also shows that production of IFN-γ cytokine is essential to activate the anti-viral and anti-tumor immune responses [3, 20, 21]. However, the major challenges while designing subunit/peptide-based vaccine is to identify a precise antigenic region or peptide which can activate a specific arm of immune system [22–25]. The ideal scenario would be to experimentally test the immune response to every potential fragment or peptide of the pathogen proteome. However, the experimental approaches are highly expensive and time-consuming.

Therefore, in order to design subunit vaccines with precision it is important to develop computational methods for the prediction of potential peptides/subunit candidates which can activate the immune responses and production of cytokines [26–29]. Recently, a number of in-silico tools have been developed for the prediction of cytokine inducing peptides such as IFNepitope [30], TNFepitope [31], IL2Pred [32], IL10Pred [33], IL6Pred [34], IL13Pred [35]. These prediction methods are utilized by researchers and experimental biologist for the prediction and designing of peptide-based vaccine candidates. In order to complement the existing tools, we have made a systematic attempt to improve our method IFNepitope. In this study, we have proposed a host-specific method IFNepitope2 for the accurate prediction of IFN-γ inducing or non-inducing peptides in human and mouse hosts. A number of composition based features and a huge amount of experimentally validated IFN-γ inducing or non-inducing peptide datasets has been incorporated in order to develop a highly accurate tool.

## Materials and Methods

### Overall Design

The workflow adapted in this study is depicted in Figure 1.

**Figure 1:**
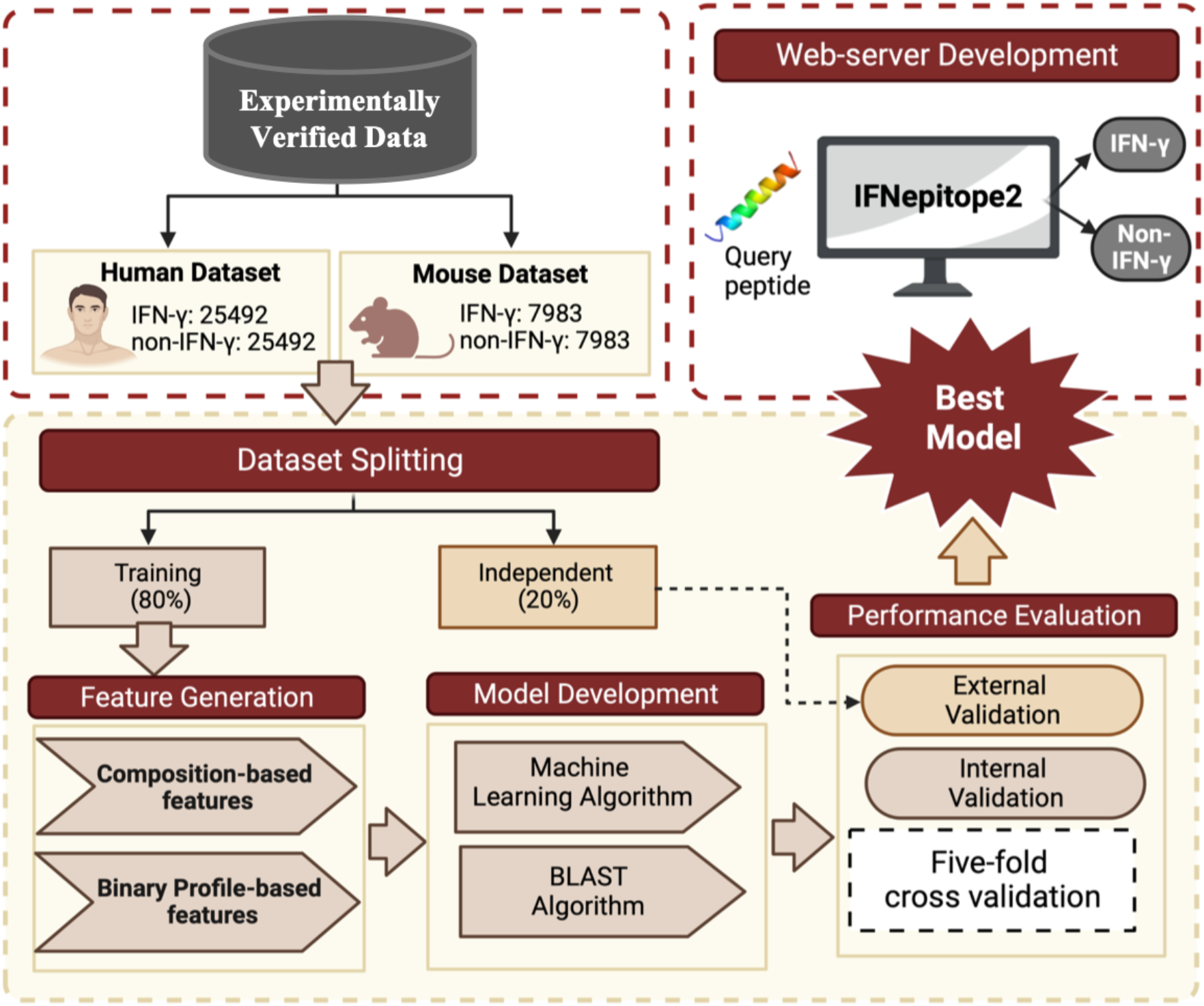
The overall architecture of the IFNepitope2, including dataset collection, feature generation, model development.

### Dataset preparation and preprocessing

In the current study, we have downloaded the experimentally validated IFN-γ inducing and non-inducing peptides from the immune epitope database (IEDB [36]). Initially, we preprocessed the downloaded file by removing the peptides having non-natural amino acids. On performing the host-based analysis, we have found that majority of the sequences belong to either human or mouse host. Therefore, we considered sequences belong to human and mouse host for further analysis. Then, we did the analysis based on length and observed that most of the sequences were lying in the length range of 8-20, hence, we selected sequences with length range 8-20 and discarded the rest. Further, we removed the redundant IFN-γ inducing and non-inducing peptides from each host. Eventually, we were left with 25492 IFN-γ inducing and 61680 non-inducing peptides belong to human host, and 7983 IFN-γ inducing and 27837 non-inducing peptides from mouse host. On further analysis, we have found out that there is high similarity in the IFN-γ inducing and non-inducing sequences in human as well as in mouse host. In order to handle that, we have removed sequences from the non-inducing peptides which either contains IFN-γ inducing peptides or differing with only one or two amino acids. We were left with 25492 IFN-γ inducing and 41092 non-inducing peptides in case of human host, and 7983 IFN-γ inducing and 16121 non-inducing peptides in mouse host. Finally, to make the balanced datasets for further consideration, we have randomly selected the equal number of sequences as of number IFN-γ inducing peptides, from the set of non-inducing peptides for each host organism. The final dataset consists of 25492 IFN-γ inducing and 25492 non-inducing peptides from human host, and 7983 IFN-γ inducing and 7983 non-inducing peptide sequences belong to mouse host.

### Training and Independent Dataset

For fair evaluation of the generated models, we have followed the standard procedure used in several studies [37–42] and divided the entire datasets in 80:20 ratio. The 80% of the dataset was used for the training purpose or internal validation and the rest 20% was kept aside for the external validation. In case of human host, the training dataset was comprised of 20394 IFN-γ inducing and non-inducing peptides where independent dataset contained 5098 IFN-γ inducing peptides and equal number of non-inducing peptides. Similarly, for mouse host, 6387 IFN-γ inducing and 6387 non-inducing peptide sequences were used in the training dataset, and 1596 IFN-γ inducing and non-inducing peptides were used as the independent dataset. Moreover, we tried to maintain the distribution of length of peptides in training and independent dataset.

### Composition Analysis

To understand the preference of amino acids in the IFN-γ inducing peptides as compare to noninducing peptides in both the hosts, we have calculated the amino acid composition of each peptide in IFN-γ inducing and non-inducing datasets. Further, we compared the average composition of each residue in IFN-γ inducing and non-inducing dataset for human as well as mouse host. We have implemented the composition-based module of Pfeature [43] to compute the amino acid composition of each peptide.

### Positional Preference Analysis

The composition analysis provides the overall preference of each amino acid in the IFN-γ inducing and non-inducing dataset, but fails to capture the preference of residues at a particular position in the peptides. In order to achieve that, we have generated the sequence logos using the two-sample logo software [44]. One of the major challenges with TSL tool was that it does not takes variable length peptides as input, therefore, it was required to fix the peptide length. The minimum length of the peptides in the dataset was 8, therefore, to generate the fixed length peptides, we have considered the 8 amino acids from the N-terminal and 8 residues from the C-terminal. Further, we joined them to generate peptides of length 16, which were used as the input to generate the sequence logos for both human and mouse datasets.

### Feature Generation

To build the prediction models with the ability to discriminate the IFN-γ inducing peptides from the non-inducing peptides, it is necessary to represent the sequences using numerical vectors. Thus, we have implemented the composition- and binary profile-based modules of Pfeature to calculate different types of features. We have computed composition-based such as amino acid composition, amphiphilic pseudo amino acid composition, atomic composition, bond composition, composition-enhanced transition distribution, conjoint triad composition, distance distribution of residues, dipeptide composition, pseudo amino acid composition, physicochemical properties composition, quasi-sequence order, residue repeats information, sequence order coupling number, Shannon-entropy of physicochemical properties, and combination of all the features. Additionally, we have also calculated four binary profiles such as binary profile eight residues from N-terminal (N8) as eight was the minimum length of the peptides, binary profile of eight residues from C-terminal (C8), combination of N8 and C8 (N8C8), and binary profile using 20 residues (NC20) where dummy residue “X” was added to make up the length 20 where peptide length was less than 20 [37]. We have used all the abovementioned features to develop the various classification models to classify the IFN-γ inducing peptides.

### Model Development

In order to build the prediction models to segregate IFN-γ inducing peptides from the noninducing peptides, we have implemented various machine learning classifiers using scikit-learn library of Python. These classifiers include decision tree (DT), random forest (RF), logistic regression (LR), K-nearest neighbor (KNN), Gaussian naïve Bayes (GNB), extra trees (ET), and support vector classifier (SVC). We have trained different classifiers using the training dataset and evaluated them on the independent dataset.

### BLAST Approach

After that, we have implemented a similarity search approach using BLAST [45], where we categories the epitopes as IFN-inducing or non-inducing based on the similarity of the sequences. In this case, we created a custom database using the makeblastdb suite of NCBI-BLAST+ version 2.2.29 and performed similarity searches using blastp suite. Using the training dataset, we built a unique database, and validation dataset sequences were checked against it. We categories the hits as IFN-inducer or non-inducer based on their similarity to the customized database. At the moment, we take only the top-hit of BLAST (i.e., if the top-hit of BLAST is against the IFN-inducer peptide, the query sequence was classified as IFN-inducing peptide or vice-versa. We run BLAST at a variety of e-value cutoffs ranging from 1e-6 to 1e+3 in order to determine the optimal e-value value score.

### Hybrid Model

We have used a hybrid strategy that combines BLAST and machine learning-based prediction approach in order to enhance the performance of prediction models. Here, we initially categories the peptide or epitope in accordance with the BLAST query. Then, we integrate ‘-0.5’ score for the incorrect or negative predictions, ‘+0.5’ score for the correct or positive predictions, and ‘0’ value if no hits were discovered. Finally, we integrate the BLAST score and the machine learning prediction score to make the final predictions.

### Cross-validation Technique

To avoid the overfitting and biasness of the generated prediction models, we have implemented the five-fold cross-validation technique on the training dataset to perform the internal validation. In this technique, the entire training dataset was first divided into five parts out of which four were used to train the model where the remaining one part was used for the testing. The same procedure was repeated five times so that each part gets the chance to be utilized for testing the model [46–48]. The final performance is evaluated by taking the mean of the performances from each iteration.

### Evaluation Metrices

In order to evaluate the performance of the models developed using various machine learning classifiers, we have used the standard performance evaluation parameters. We have computed the threshold-dependent and threshold-independent parameters. Threshold-dependent parameters included sensitivity, specificity, accuracy, F1-score, Kappa, and Matthews correlation coefficients. The equations for the threshold-dependent metrices are represented in equation 1–5. On the other hand, area-under the receiver operating characteristics curve was used as threshold-independent parameter.

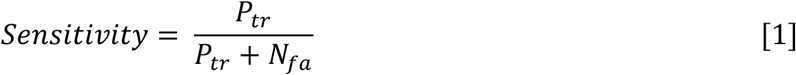

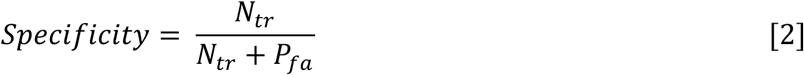

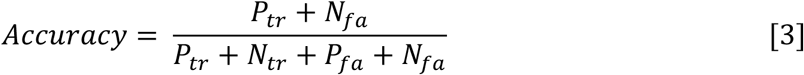

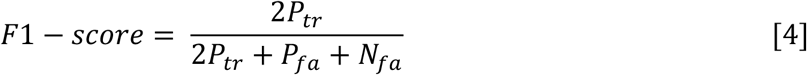

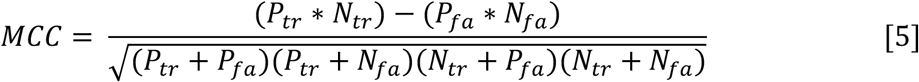

Where, *P_tr_* is true positive, *N_tr_* is true negative, *P_fa_* is false positive, and *N_fa_* is false negative.

## Results

### Composition and conservation analysis

At first, we computed amino acid composition of IFN-inducing and non-inducing peptides using the human and mouse datasets. Figure 2A and 2C shows the average amino acid for human and mouse dataset. As shown in Figure 2A, the average amino acid composition of residues (K, P and N) is higher in IFN-γ inducing peptides in the human dataset. However, the average composition of residues such as A, G, and P is higher in mouse IFN-inducing peptides (See Figure 2C). In the case of human host IFN-inducing epitopes, residue ‘K’ are highly conserved at the majority of positions, whereas ‘P’ is preferred at the 6^th^, 7^th^, 15^th^ and 16^th^ positions (See Figure 2B). In IFN-inducing epitopes of mouse host, residues ‘A’ and ‘P’ highly dominated at most of the positions (See Figure 2D).

**Figure 2:**
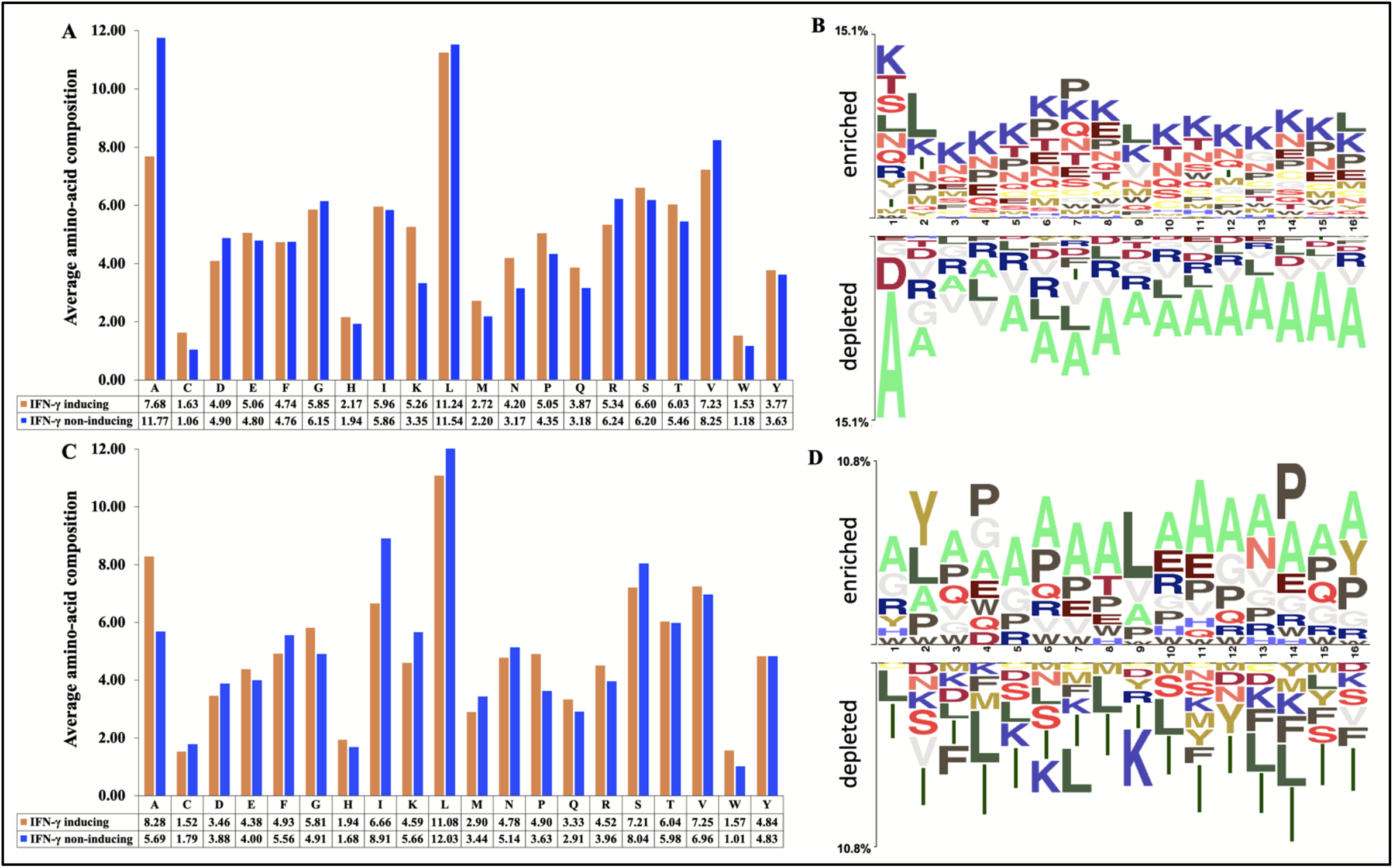
Average amino-acid composition and two sample logo of IFN-γ inducing and non-inducing peptide, (A and B) Human dataset (C and D) Mouse dataset.

### Performance of Machine Learning Models

Various prediction models were developed using seven classifiers such as DT, RF, ET, GNB, KNN, LR and SVC for both human and mouse datasets. We have used Pfeature standalone package to compute binary and composition based features. Then we evaluated the performance on different features as well as combining all the features on training and independent datasets.

### Performance of Composition-based Models

We have developed various composition-based features using human dataset. We observed that ET classifier achieved highest performance for most of the feature types. In Table 1, we have provided the performance of all the features obtained using ET classifier. The complete results of all the classifiers for the different features are given in Supplementary Table S1. As shown in Table 1, the di-peptide composition-based feature achieved highest AUROC of (0.90 and 0.89) and MCC (0.63 and 0.61) on training and independent dataset. ALL_COMB based-features achieved comparable performance in terms of AUROC (0.89 and 0.88) and MCC (0.61 and 0.60) on training and independent datasets. Similarly, AAC, APAAC and PAAC able to achieve equivalent performances with an AUROC of 0.88 on training and 0.79 on independent dataset, with balanced sensitivity and specificity. PCP, DDR, QSO, SPC and CETD features perform quite well with an AUROC > 0.80 on training and validation datasets. As shown in Supplementary Table S1, RF and SVC classifier also achieved comparable performances on independent dataset. We found RF achieved maximum AUROC of 0.87 using ALL_COMP features, and SVC achieved maximum AUROC of 0.8 on DPC-based features on training and validation datasets, respectively (See Supplementary Table S1).

**Table 1:** Performance metrices of extra tree (ET) classifier build using various composition-based features on training and independent dataset of human host.

| Feature Type | Training Dataset |  |  |  |  |  | Independent Dataset |  |  |  |  |  |
| --- | --- | --- | --- | --- | --- | --- | --- | --- | --- | --- | --- | --- |
|  | Sens | Spec | ACC | AUROC | F1 | MCC | Sens | Spec | ACC | AUROC | F1 | MCC |
| AAC | 80.35 | 79.43 | 79.89 | 0.88 | 0.80 | 0.60 | 79.25 | 78.93 | 79.09 | 0.87 | 0.79 | 0.58 |
| DPC | 80.73 | 82.29 | 81.51 | 0.90 | 0.82 | 0.63 | 81.56 | 79.38 | 80.47 | 0.89 | 0.81 | 0.61 |
| BTC | 64.29 | 64.86 | 64.58 | 0.68 | 0.65 | 0.29 | 65.81 | 62.38 | 64.09 | 0.68 | 0.65 | 0.28 |
| ATC | 58.70 | 59.97 | 59.34 | 0.63 | 0.59 | 0.19 | 58.55 | 60.26 | 59.41 | 0.63 | 0.59 | 0.19 |
| PCP | 76.50 | 77.05 | 76.77 | 0.84 | 0.77 | 0.54 | 75.44 | 77.27 | 76.35 | 0.84 | 0.76 | 0.53 |
| DDR | 76.08 | 74.59 | 75.33 | 0.84 | 0.76 | 0.51 | 75.21 | 74.30 | 74.76 | 0.83 | 0.75 | 0.50 |
| RRI | 67.49 | 68.62 | 68.05 | 0.74 | 0.68 | 0.36 | 65.14 | 67.71 | 66.43 | 0.72 | 0.66 | 0.33 |
| PAAC | 80.48 | 79.59 | 80.03 | 0.88 | 0.80 | 0.60 | 79.48 | 78.31 | 78.89 | 0.87 | 0.79 | 0.58 |
| APAAC | 80.04 | 80.64 | 80.34 | 0.88 | 0.80 | 0.61 | 78.72 | 79.36 | 79.04 | 0.87 | 0.79 | 0.58 |
| SOC | 51.49 | 50.17 | 50.83 | 0.52 | 0.51 | 0.02 | 51.59 | 50.63 | 51.11 | 0.51 | 0.51 | 0.02 |
| SPC | 76.35 | 75.97 | 76.16 | 0.84 | 0.76 | 0.52 | 76.13 | 76.68 | 76.40 | 0.83 | 0.76 | 0.53 |
| CTC | 67.77 | 68.66 | 68.21 | 0.76 | 0.68 | 0.36 | 68.03 | 67.05 | 67.54 | 0.74 | 0.68 | 0.35 |
| CETD | 76.90 | 76.48 | 76.69 | 0.85 | 0.77 | 0.53 | 76.50 | 76.83 | 76.67 | 0.84 | 0.77 | 0.53 |
| QSO | 72.90 | 73.24 | 73.07 | 0.81 | 0.73 | 0.46 | 71.17 | 72.79 | 71.98 | 0.80 | 0.72 | 0.44 |
| ALL_COMB | 80.64 | 80.71 | 80.67 | 0.89 | 0.81 | 0.61 | 79.87 | 80.05 | 79.96 | 0.88 | 0.80 | 0.60 |

\*Sens: Sensitivity; Spec: Specificity; Acc: Accuracy; AUROC: Area Under the Receiver Operating Characteristics curve; MCC: Matthews Correlation Coefficient; AAC: Amino Acid Composition; DPC: Di-peptide Composition; ATC: Atomic Composition; APAAC: Amphiphilic Pseudo Amino Acid Composition; BTC: Bond Composition; CETD: Composition-Enhanced Transition Distribution; CTC: Conjoint Triad Descriptors; DDR: Distance Distribution of Residues; PAAC: Pseudo Amino Acid Composition; PCP: Physico-Chemical Properties composition; QSO: Quasi-Sequence Order; RRI: Residue Repeat Information; SOC: Sequence Order Coupling number; SEP: Shannon-Entropy of Peptide; SER: Shannon-Entropy of Residues; SPC: Shannon-entropy of Physico-Chemical properties; ALL\_COMP: Combination of All Composition based features

In order to identify IFN-inducing epitopes in mouse host, we have developed similar prediction models for mouse datasets. We observed that in case of mouse host, ET based classifier perform well on most of the feature, hence we have provided performance of ET in Table 2. The overall results of other classifiers are provided in Supplementary Table S2. As shown in Table 2, dipeptide based features performed best among the other feature type. We obtained a highest AUROC of 0.88 and 0.83; MCC of 0.60 and 0.51 on training and independent datasets. ALL_COMB features also performed well on both training and validation datasets with an AUROC of 0.83 and 0.82; MCC of 0.49 and 0.48, respectively. Whereas, AAC, APAAC and PAAC performed equivalent with an AUROC of 0.80 and 0.79 on training and validation datasets. Similarly, we observed that RF based models also perform well on independent dataset. As shown in Supplementary Table S2, for AAC and DPC based features RF achieved maximum AUROC of (0.78 and 0.85) and AUROC of (0.77 and 0.83) on training and independent dataset.

**Table 2:** Performance metrices of extra tree (ET) build using various composition-based features on training and independent dataset of mouse host.

| Feature Type | Training Dataset |  |  |  |  |  | Independent Dataset |  |  |  |  |  |
| --- | --- | --- | --- | --- | --- | --- | --- | --- | --- | --- | --- | --- |
|  | Sens | Spec | ACC | AUROC | F1 | MCC | Sens | Spec | ACC | AUROC | F1 | MCC |
| AAC | 72.23 | 73.18 | 72.70 | 0.80 | 0.73 | 0.45 | 71.24 | 72.37 | 71.81 | 0.79 | 0.72 | 0.44 |
| DPC | 79.77 | 80.21 | 79.99 | 0.88 | 0.80 | 0.60 | 76.00 | 75.13 | 75.56 | 0.83 | 0.76 | 0.51 |
| BTC | 55.38 | 53.01 | 54.20 | 0.56 | 0.55 | 0.08 | 61.53 | 50.06 | 55.80 | 0.57 | 0.58 | 0.12 |
| ATC | 54.94 | 56.69 | 55.82 | 0.58 | 0.55 | 0.12 | 57.52 | 56.20 | 56.86 | 0.59 | 0.57 | 0.14 |
| PCP | 65.57 | 67.95 | 66.76 | 0.74 | 0.66 | 0.34 | 66.48 | 68.80 | 67.64 | 0.73 | 0.67 | 0.35 |
| DDR | 69.70 | 68.53 | 69.12 | 0.77 | 0.69 | 0.38 | 68.48 | 67.79 | 68.14 | 0.75 | 0.68 | 0.36 |
| **RRI** | 67.78 | 68.31 | 68.04 | 0.75 | 0.68 | 0.36 | 67.54 | 67.67 | 67.61 | 0.73 | 0.68 | 0.35 |
| **PAAC** | 72.84 | 73.57 | 73.20 | 0.81 | 0.73 | 0.46 | 71.81 | 72.18 | 71.99 | 0.79 | 0.72 | 0.44 |
| **APAAC** | 73.04 | 72.87 | 72.95 | 0.80 | 0.73 | 0.46 | 72.49 | 70.99 | 71.74 | 0.79 | 0.72 | 0.44 |
| **SOC** | 50.99 | 50.98 | 50.99 | 0.52 | 0.51 | 0.02 | 52.32 | 51.19 | 51.75 | 0.53 | 0.52 | 0.04 |
| **SPC** | 67.02 | 65.86 | 66.44 | 0.72 | 0.66 | 0.33 | 67.29 | 65.28 | 66.29 | 0.71 | 0.66 | 0.35 |
| **CTC** | 69.16 | 67.50 | 68.33 | 0.75 | 0.69 | 0.37 | 67.92 | 62.97 | 65.45 | 0.71 | 0.66 | 0.31 |
| **CETD** | 67.36 | 69.06 | 68.21 | 0.76 | 0.68 | 0.36 | 68.17 | 67.92 | 68.05 | 0.75 | 0.68 | 0.36 |
| **QSO** | 70.21 | 70.85 | 70.53 | 0.78 | 0.70 | 0.41 | 71.05 | 69.61 | 70.33 | 0.76 | 0.71 | 0.41 |
| **ALL_COMB** | 73.82 | 75.48 | 74.65 | 0.83 | 0.74 | 0.49 | 73.94 | 73.87 | 73.90 | 0.82 | 0.74 | 0.48 |

\*Sens: Sensitivity; Spec: Specificity; Acc: Accuracy; AUROC: Area Under the Receiver Operating Characteristics curve; MCC: Matthews Correlation Coefficient; AAC: Amino Acid Composition; DPC: Di-peptide Composition; ATC: Atomic Composition; APAAC: Amphiphilic Pseudo Amino Acid Composition; BTC: Bond Composition; CETD: Composition-Enhanced Transition Distribution; CTD: Conjoint Triad Descriptors; DDR: Distance Distribution of Residues; PAAC: Pseudo Amino Acid Composition; PCP: Physico-Chemical Properties composition; QSO: Quasi-Sequence Order; RRI: Residue Repeat Information; SOC: Sequence Order Coupling number; SEP: Shannon-Entropy of Peptide; SER: Shannon-Entropy of Residues; SPC: Shannon-entropy of Physico-Chemical properties; ALL\_COMP: Combination of All Composition based features

### Performance of Binary Profile-based Models

In addition, we have generated binary profile-based features and computed performances on binary profile of eight residues from N-terminal (N8), binary profile of eight residues from C-terminal (C8), combination of N8 and C8 (N8C8), and binary profile using 20 residues (NC20). As shown in the Table 3, ET classifier perform best among the other classifiers using NC20 binary profile-based features. It achieves maximum AUROC of 0.86 and MCC of 0.56 on both training and independent datasets. Similarly, N8C8 type features also perform well with an AUROC of 0.80 training and 0.79 independent datasets (See Supplementary Table S3). On the other side, in the case of mouse dataset we were able to achieve maximum AUROC of 0.76 and MCC of 0.37 on independent dataset using NC20 binary profile-based features (See Table 3). In addition, we have obtained comparable performance of AUROC 0.70 and 0.71 using SVC classifier and N8C8 features, on training and independent dataset. The complete results of all the other classifiers for human and mouse hosts is provided in Supplementary Table S3. Of note, we observed that for both human and mouse hosts the di-peptide composition-based features performed best on independent datasets (See Table 1 and Table 2).

**Table 3:** Performance of extra tree (ET) classifier-based models build using binary profile-based features on training and independent dataset of human and mouse host.

| Feature Type | Training Dataset |  |  |  |  |  | Independent Dataset |  |  |  |  |  |
| --- | --- | --- | --- | --- | --- | --- | --- | --- | --- | --- | --- | --- |
|  | Human Dataset |  |  |  |  |  |  |  |  |  |  |  |
|  | Sens | Spec | ACC | AUROC | F1 | MCC | Sens | Spec | ACC | AUROC | F1 | MCC |

| N8 | 63.52 | 64.30 | 63.91 | 0.70 | 0.64 | 0.28 | 62.91 | 64.10 | 63.51 | 0.70 | 0.63 | 0.27 |
| --- | --- | --- | --- | --- | --- | --- | --- | --- | --- | --- | --- | --- |
| C8 | 63.84 | 62.13 | 62.98 | 0.69 | 0.63 | 0.26 | 63.85 | 61.22 | 62.53 | 0.68 | 0.63 | 0.25 |
| N8C8 | 71.37 | 69.60 | 70.49 | 0.78 | 0.71 | 0.41 | 72.17 | 69.32 | 70.74 | 0.78 | 0.71 | 0.42 |
| NC20 | 77.73 | 78.20 | 77.96 | 0.86 | 0.78 | 0.56 | 78.01 | 78.07 | 78.04 | 0.86 | 0.78 | 0.56 |
| Feature Type | Mouse Dataset |  |  |  |  |  |  |  |  |  |  |  |
|  | Sens | Spec | ACC | AUROC | F1 | MCC | Sens | Spec | ACC | AUROC | F1 | MCC |
| N8 | 62.85 | 60.87 | 61.86 | 0.68 | 0.62 | 0.24 | 61.40 | 62.03 | 61.72 | 0.68 | 0.62 | 0.23 |
| C8 | 63.10 | 62.30 | 62.70 | 0.69 | 0.63 | 0.25 | 64.04 | 61.40 | 62.72 | 0.68 | 0.63 | 0.25 |
| N8C8 | 64.30 | 65.67 | 64.98 | 0.71 | 0.65 | 0.30 | 64.04 | 65.29 | 64.66 | 0.71 | 0.64 | 0.29 |
| NC20 | 69.88 | 68.95 | 69.41 | 0.76 | 0.70 | 0.39 | 68.11 | 68.92 | 68.52 | 0.76 | 0.68 | 0.37 |

\*Sens: Sensitivity; Spec: Specificity; Acc: Accuracy; AUROC: Area Under the Receiver Operating Characteristics curve; MCC: Matthews Correlation Coefficient;

### Performance of Hybrid Models

In order to predict IFN-inducing and non-inducing peptides, we have built a hybrid model by combining BLAST and machine learning approach. Here, we have combined di-peptide based predictions and BLAST similarity scores. As we observed di-peptide composition-based features performed best among the other features, in case of both human and mouse hosts (See Table 1 and Table 2). So, we generated hybrid models using di-peptide based features and BLAST alignment score. As shown in Table 4, we have calculated the performances at various e-value cutoffs. In the case of human models, we achieve maximum performance at e-value (evalue (1.00E-06) with AUC of 0.92 and MCC of 0.70 on training datasets and AUC of 0.90 and MCC of 0.66 on independent datasets. In case of mouse dataset, we obtained maximum performance at e-value (1.00E-06) with AUC of 0.89 and MCC of 0.62 on training datasets and AUC 0.85 and MCC 0.55 on independent datasets (See Table 4). The complete results of hybrid models are available in Supplementary Table S4.

**Table 4:** Performance of hybrid model on different E-values, build by integrating BLAST and machine learning approach on training and independent datasets for human and mouse host.

| E-value | Training Dataset |  |  |  |  |  | Independent Dataset |  |  |  |  |  |
| --- | --- | --- | --- | --- | --- | --- | --- | --- | --- | --- | --- | --- |
|  | Human Dataset |  |  |  |  |  |  |  |  |  |  |  |
|  | Sens | Spec | ACC | AUC | F1 | MCC | Sens | Spec | ACC | AUC | F1 | MCC |
| 1.00E-06 | 84.54 | 85.24 | 84.89 | 0.92 | 0.85 | 0.70 | 83.31 | 82.88 | 83.09 | 0.90 | 0.83 | 0.66 |
| 1.00E-05 | 84.29 | 85.22 | 84.76 | 0.92 | 0.85 | 0.70 | 83.35 | 82.68 | 83.01 | 0.90 | 0.83 | 0.66 |
| 1.00E-04 | 84.26 | 85.10 | 84.68 | 0.91 | 0.85 | 0.69 | 83.41 | 82.62 | 83.01 | 0.89 | 0.83 | 0.66 |

| 1.00E-03 | 84.15 | 84.90 | 84.53 | 0.91 | 0.85 | 0.69 | 83.54 | 82.23 | 82.89 | 0.89 | 0.83 | 0.66 |
| --- | --- | --- | --- | --- | --- | --- | --- | --- | --- | --- | --- | --- |
| 1.00E-02 | 84.37 | 84.75 | 84.56 | 0.90 | 0.85 | 0.69 | 83.54 | 82.29 | 82.92 | 0.88 | 0.83 | 0.66 |
| 1.00E-01 | 84.52 | 83.91 | 84.22 | 0.90 | 0.84 | 0.68 | 84.13 | 81.76 | 82.94 | 0.88 | 0.83 | 0.66 |
| 1.00E+00 | 84.51 | 83.22 | 83.87 | 0.89 | 0.84 | 0.68 | 83.76 | 81.40 | 82.58 | 0.88 | 0.83 | 0.65 |
| 1.00E+01 | 83.05 | 83.29 | 83.17 | 0.89 | 0.83 | 0.66 | 82.68 | 82.07 | 82.38 | 0.88 | 0.82 | 0.65 |
| 1.00E+02 | 75.40 | 75.04 | 75.22 | 0.88 | 0.75 | 0.50 | 76.44 | 76.34 | 76.39 | 0.87 | 0.76 | 0.53 |
| 1.00E+03 | 77.71 | 77.34 | 77.53 | 0.89 | 0.78 | 0.55 | 77.76 | 77.64 | 77.70 | 0.88 | 0.78 | 0.55 |
| E-value | Mouse Dataset |  |  |  |  |  |  |  |  |  |  |  |
|  | Sens | Spec | ACC | AUC | F1 | MCC | Sens | Spec | ACC | AUC | F1 | MCC |
| 1.00E-06 | 80.40 | 81.31 | 80.85 | 0.89 | 0.81 | 0.62 | 76.75 | 77.82 | 77.29 | 0.85 | 0.77 | 0.55 |
| 1.00E-05 | 80.41 | 81.04 | 80.73 | 0.89 | 0.81 | 0.62 | 76.38 | 77.44 | 76.91 | 0.85 | 0.77 | 0.54 |
| 1.00E-04 | 81.82 | 79.38 | 80.60 | 0.88 | 0.81 | 0.61 | 77.95 | 76.19 | 77.07 | 0.84 | 0.77 | 0.54 |
| 1.00E-03 | 80.48 | 81.15 | 80.81 | 0.88 | 0.81 | 0.62 | 75.50 | 78.20 | 76.85 | 0.84 | 0.77 | 0.54 |
| 1.00E-02 | 79.80 | 80.85 | 80.33 | 0.88 | 0.80 | 0.61 | 76.57 | 78.51 | 77.54 | 0.84 | 0.77 | 0.55 |
| 1.00E-01 | 81.43 | 80.04 | 80.73 | 0.88 | 0.81 | 0.62 | 76.88 | 77.19 | 77.04 | 0.84 | 0.77 | 0.54 |
| 1.00E+00 | 80.74 | 80.21 | 80.48 | 0.88 | 0.81 | 0.61 | 76.94 | 77.26 | 77.10 | 0.84 | 0.77 | 0.54 |
| 1.00E+01 | 80.77 | 79.85 | 80.31 | 0.87 | 0.80 | 0.61 | 76.94 | 77.51 | 77.22 | 0.85 | 0.77 | 0.54 |
| 1.00E+02 | 75.92 | 76.47 | 76.19 | 0.87 | 0.76 | 0.52 | 81.08 | 74.37 | 77.73 | 0.77 | 0.78 | 0.56 |
| 1.00E+03 | 75.89 | 75.31 | 75.60 | 0.81 | 0.76 | 0.51 | 73.00 | 80.70 | 76.85 | 0.85 | 0.76 | 0.54 |

\*Sens: Sensitivity; Spec: Specificity; Acc: Accuracy; AUROC: Area Under the Receiver Operating Characteristics curve; MCC: Matthews Correlation Coefficient;

### Web-server Development

For the prediction of IFN-inducing and non-inducing epitopes using sequence information, we have created a web service called “IFNepitope2”. The server is developed using HTML, JAVA, and PHP scripts and is compatible with a variety of gadgets, including laptops, iPhones, phones, etc. We have incorporated the hybrid models (DPC + BLAST) at the back-end of the webserver. IFNepitope2 includes three major modules: ‘Predict’, ‘Scan’, and ‘Design’. The ‘Predict’ module classify the IFN-inducing and non-inducing peptides; and allow the user to provide peptide sequences in FASTA format. ‘Scan’ module provides the facility to the user to map/scan the IFN-gamma inducing fragments in the protein sequences. ‘Design’ module facilitates the user to generate the possible mutants of the query sequence and predict the mutated sequence have the capacity to secrete IFN-gamma inducing/non-inducing peptides. Moreover, the training and validation datasets were available on the download page of the webserver. The webserver is available at https://webs.iiitd.edu.in/raghava/ifnepitope2/ link.

### Comparison with existing tool

It is essential to compare the performance of newly developed method, with the existing method in order to validate it. In the past there is only method for the prediction of IFN-gamma inducing peptides [30]. Hence, we compare our method with the existing one on same independent dataset to provide unbiased evaluation. The performance measures such as sensitivity, specificity, AUROC, accuracy, F1, and MCC were computed to compare the performance of both the methods (See Supplementary Table S5). AUROC plots show that IFNepitope2 outperforms the existing method on both human and mouse hosts (See Figure 3).

**Figure 3:**
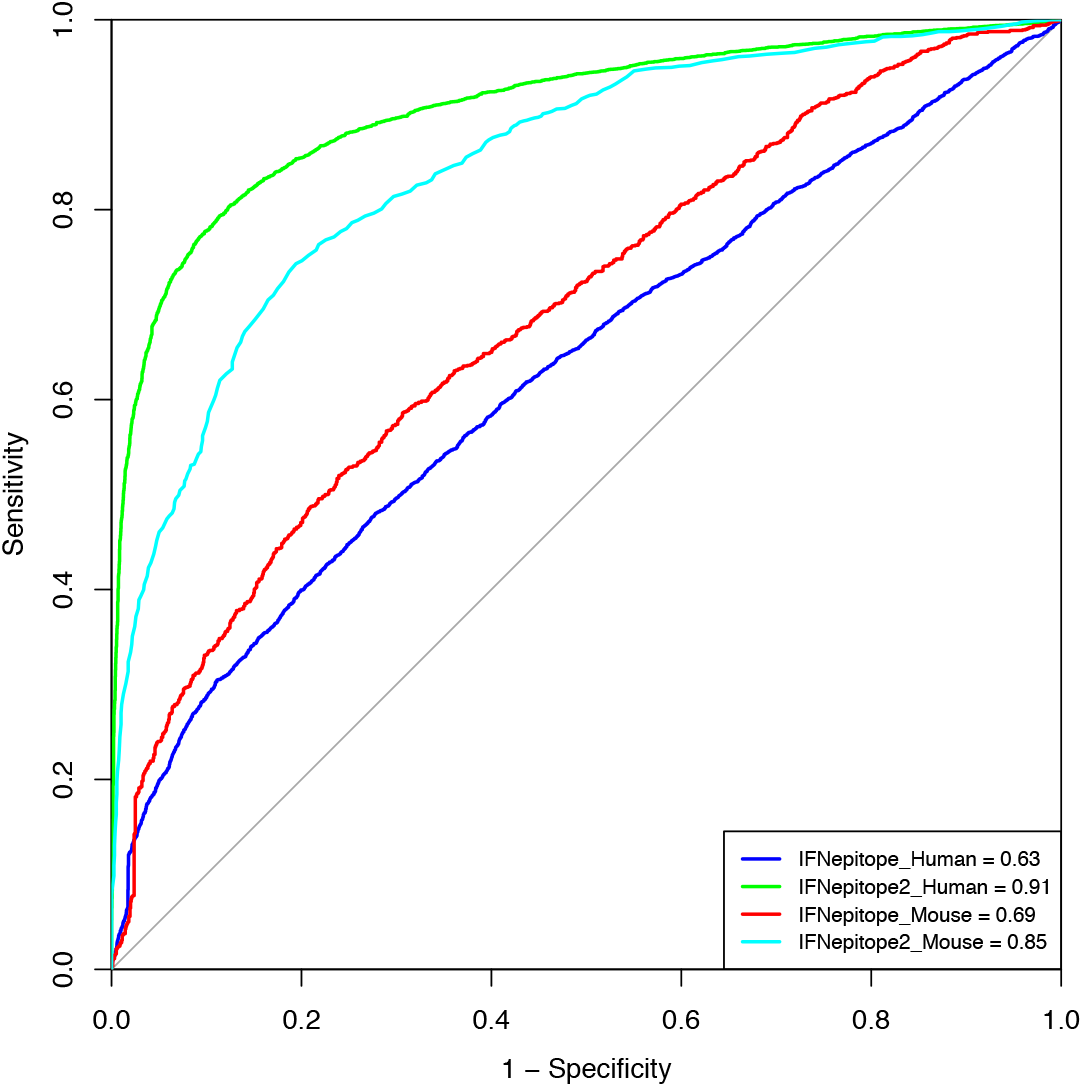
Comparison of performance in terms of AUROC between IFNepitope and IFNepitope2 on the independent dataset for human and mouse host.

## Discussion & Conclusion

Traditional vaccinations are made up of whole pathogens that have been destroyed or weekend so they can no longer spread illness. Whole-pathogen vaccines can stimulate protective immune responses against pathogens and infections [49, 50]. However, it is not necessary the entire protein is effective in targeting the pathogen or eliciting the desired immune response. Hence, subunit/peptide-based vaccines are developed in which a small portion of pathogen protein is utilized in order to generate the vaccine candidates against [24, 51–56]. While designing a subunit-vaccines it is necessary to check the peptide/antigens have the capacity to effectively stimulate the immune system and generate specific type of cytokines to send signal to the immune system to perform inflammatory responses against the pathogens and cancerous cells. IFN-γ is one of most important cytokines which generate anti-tumor, anti-viral and proinflammatory responses against pathogens [57–59]. Hence, it is necessary to check the IFN-inducing potential subunit vaccine candidates. In order to help the researchers and experimental biologist we have developed a highly accurate tool for the prediction and designing of IFN-γ inducing peptides.

In the current study, we have used experimentally validated IFN-γ inducing and non-inducing peptides from IEDB resource. We have developed host-specific prediction models using human and mouse datasets. At first, we performed composition and positional preference analysis using human and mouse datasets. We observed that the average composition of ‘Lysine’, ‘Proline’ and ‘Asparagine’ amino acid is very high for human IFN-γ inducing peptides. While the composition of ‘Alanine’ and ‘Proline’ is higher in the case of mouse IFN-γ inducing peptides. Similarly, TSL plot reveals that ‘Lysine’ residue is highly preferred at most of the position in case of human and ‘Alanine’ is preferred in case of mouse dataset. We generated features binary-profile and composition-based features using Pfeature standalone package. We have developed various machine learning classifiers using number of features for both human and mouse hosts. Our hybrid model is able to achieve a maximum AUROC of (0.92 and 0.90) for human host and AUROC of (0.89 and 0.85) for mouse host on training and independent datasets, respectively. We have used the best models for both the host to generate the web-server (IFNepitope2) and standalone package. Moreover, we also compared the performance of our models with the existing tool. Our tool ‘IFNepitope2’ outperform the existing tool on independent validation for both human and mouse hosts. We anticipate this newly developed method is validated on huge amount of experimentally validated dataset and highly accurate then the existing method.

## Authors’ contributions

AD and GPSR collected and processed the datasets. AD, SP and GPSR implemented the algorithms and developed the prediction models. AD, SP and GPSR analysed the results. AD and SP created the web server. AD, SP and GPSR penned the manuscript. GPSR conceived and coordinated the project. All authors have read and approved the final manuscript.

## Acknowledgements

Authors are thankful to the Department of Bio-Technology (DBT) and Department of Science and Technology (DST-INSPIRE) for fellowships and the financial support and Department of Computational Biology, IIITD New Delhi for infrastructure and facilities.

